# Leveraging multiple data types for improved compound-kinase bioactivity prediction

**DOI:** 10.1101/2024.03.07.583951

**Authors:** Ryan Theisen, Tianduanyi Wang, Balaguru Ravikumar, Rayees Rahman, Anna Cichońska

## Abstract

Machine learning methods offer time- and cost-effective means for identifying novel chemical matter as well as guiding experimental efforts to map enormous compound-kinase interaction spaces. However, considerable challenges for compound-kinase interaction modeling arise from the heterogeneity of available bioactivity readouts, including single-dose compound profiling results, such as percentage inhibition, and multi-dose-response results, such as IC_50_ . Standard activity prediction approaches utilize only dose-response data in the model training, disregarding a substantial portion of available information contained in single-dose measurements. Here, we propose a novel machine learning methodology for compound-kinase activity prediction that leverages both single-dose and dose-response data. Our two-stage model first learns a mapping between single-dose and dose-response bioactivity readouts, and then generates proxy dose-response activity labels for compounds that have only been tested in single-dose assays. The predictions from the first-stage model are then integrated with experimentally measured dose-response activities to model compound-kinase binding based on chemical structures and kinase features. We demonstrate that our two-stage approach yields accurate activity predictions and significantly improves model performance compared to training solely on dose-response labels, particularly in the most practical and challenging scenarios of predicting activities for new compounds and new compound scaffolds. This superior performance is consistent across five evaluated machine learning methods, including traditional models such as random forest and kernel learning, as well as deep learning-based approaches. Using the best performing model, we carried out extensive experimental profiling on a total of 347 selected compound-kinase pairs, achieving a high hit rate of 40% and a negative predictive value of 78%. We show that these rates can be improved further by incorporating model uncertainty estimates into the compound selection process. By integrating multiple activity data types, we demonstrate that our approach holds promise for facilitating the development of training activity datasets in a more efficient and cost-effective way.

## 1. Introduction

The enormous size of the kinase inhibitor chemical space poses a considerable challenge for traditional experimental approaches to map compound-kinase interaction spaces, highlighting the need for alternative strategies that can expedite the kinase inhibitor discovery process. Beyond traditional molecular docking approaches, machine learning methods have emerged as promising tools in this context, offering time- and cost-effective means to navigate the kinase chemical space. In fact, several new models for compound-kinase binding prediction are introduced every month [CCAS^+^15, CRA^+^21, DQJ^+^22, DSSGP22]. They differ in the learning algorithm used, such as simple k-nearest neighbor regression [BHS^+^21], decision trees [TAA^+^22], kernel learning [MM12, NPC16, CRP^+^17, CPS^+^18] and deep learning methods [BHS^+^21, O18, KZEK23, LLP23, SSB^+^23], as well as compound and protein descriptors, including compound SMILES and graphs [DTME20], protein amino acid sequences [BHS^+^21, KZEK23] and, lately, more complex 3D structure-based features [KZK^+^23, PHL^+^23, LKN^+^23, LTZ^+^23] and embeddings from pretrained large language models [SSB^+^23]. Most recent methods modeling compound-kinase activities learn from the descriptors of both compounds and kinases, and are referred to as proteochemometric models. For example, the BiMCA model is based on a bimodal neural network that incorporates convolutional and attention layers, using text sequences of SMILES for compounds and amino acids for kinases [BHS^+^21]. On the other hand, ConPLex, another deep learning model, predicts compound-kinase activities using compound ECFP4 fingerprints, with kinase features derived from a pretrained ProtBert language model [EHD^+^21]. Furthermore, ConPLex employs a contrastive learning stage, training the model to predict activities while simultaneously learning to differentiate real drugs from synthetically generated decoys [SSB^+^23].

Although these machine learning methods have demonstrated strong performance within their respective evaluation scenarios, the available bioactivity datasets for model training are very heterogeneous, comprising a variety of data types, and thus pose significant challenges for compound-kinase interaction modeling. Specifically, compound activity against a kinase is typically determined either from a single dose of compound (e.g., percentage inhibition or activity readouts) or more comprehensive and costly dose-response profiling (e.g., dissociation constant K_d_, inhibition constant K_i_, or half-maximal inhibitory concentration IC_50_ readouts). Conventional approaches modeling compound-kinase activity rely on dose-response data only, thereby ignoring a substantial portion of the available information. The neglect of point-of-concentration (POC) measurements is particularly noteworthy given the prevalence of such data in public databases. For example, approximately 40% of all kinase activity data in ChEMBL bioactivity database [MGB^+^19] consists of compounds for which only POC activities were measured. This large pool of compounds with POC data has yet to be utilized by current activity modeling approaches, therefore limiting compound-kinase activity training spaces and potentially overlooking valuable chemical matter.

To address these limitations, we developed a two-stage machine learning methodology for compound-kinase activity prediction, integrating both single-dose and dose-response experimental readouts. In the first stage, we use a random forest model to learn a mapping from POC to dose-response activity values. This model is then employed to generate proxy dose-response activity labels for compounds with only POC measurements, thereby expanding the available dose-response training dataset. Predictions from the first-stage model, combined with experimentally measured dose-response activities, are used to predict compound-kinase binding affinities based on chemical structures and kinase features (Fig. 1).

**Figure 1.**
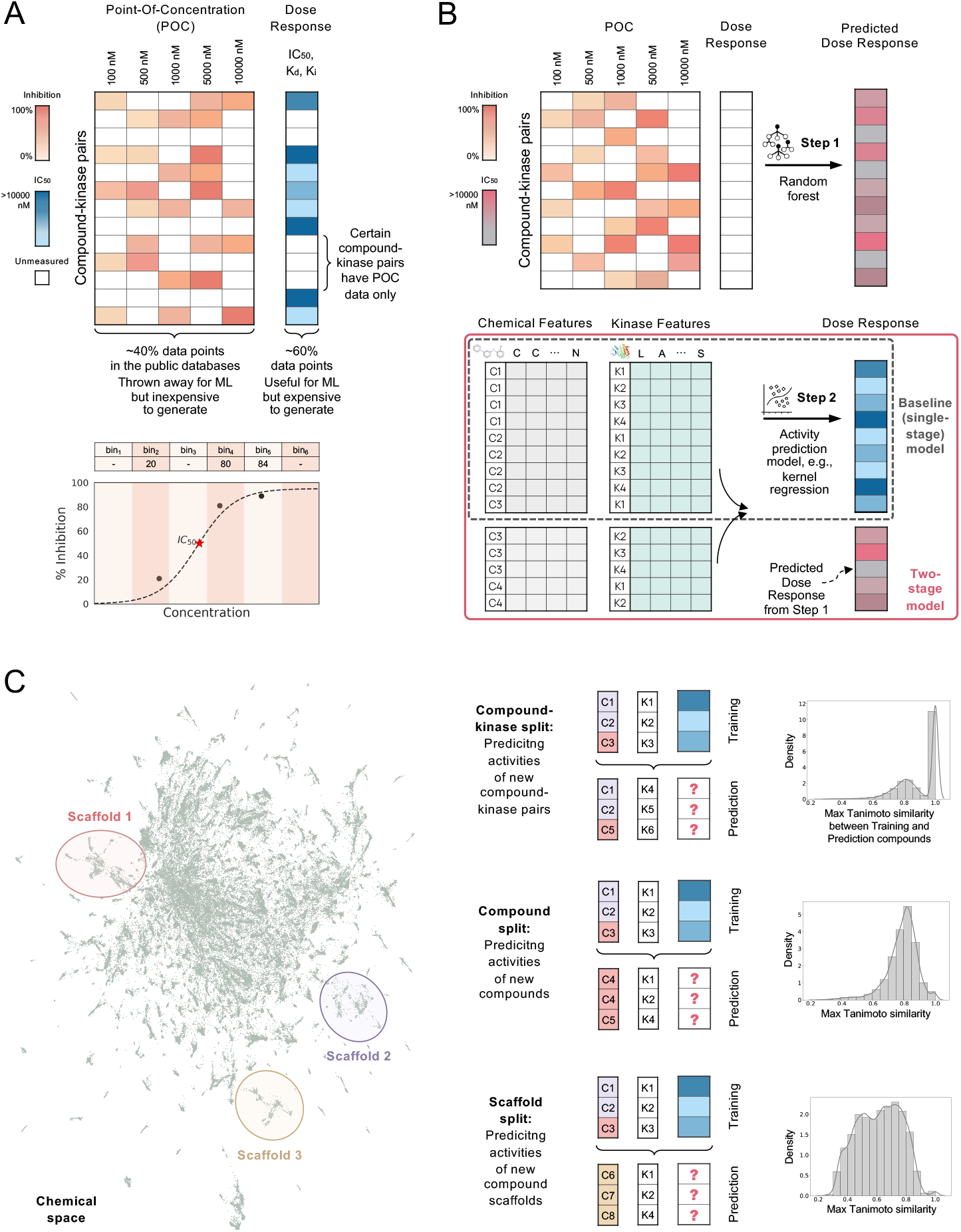
Schematic overview of the bioactivity data integration methodology for compound-kinase binding prediction. **(A)** Approximately 40% of kinase activity data in public databases comprises point-of-concentration (POC) readouts, such as percentage inhibition. These data points are relatively inexpensive to generate compared to dose-response measurements such as IC_50_, but they are typically ignored when training compound activity prediction models. **(B)** Here, we present a novel two-stage framework that integrates POC readouts with dose-response data to improve activity prediction model performance. First, a random forest model is employed to learn the mapping between POC and dose-response measurements. Subsequently, proxy dose-response labels are generated and combined with experimentally determined ones in a second-stage model. This model predicts compound-kinase activities using chemical and kinase features. **(C)** A schematic representation of the three prediction scenarios considered in this study. 1. Compound-kinase split: a prediction scenario aimed at filling in missing kinase activities for compounds that may be present during model training. 2. Compound split: a prediction scenario to infer activities of a compound that is not explicitly observed during training but may have close analogs present in the training data. 3. Scaffold split: a prediction scenario to predict activities of a series of compounds with a core scaffold not observed during training.

We demonstrate that our approach enables the exploration of a more extensive chemical space and enhances the accuracy of compound-kinase interaction predictions across various learning algorithms used in the second stage, ranging from random forest to more sophisticated kernel and deep learning methods. The most notable improvement in performance is particularly evident in the most practical and challenging scenarios of predicting the activities of new potential kinase inhibitors and kinase inhibitor scaffolds. We then use our top-performing model to screen a large purchasable compound library against 13 kinase targets and experimentally measure 297 of the most promising, previously untested compound-kinase interactions, achieving a hit rate of 40%, notably higher than those typically reported in virtual screening studies [ZCS^+^13, LWB^+^19]. Additionally, we provide a practical guide to obtaining uncertainty estimates for kernel-based activity predictions. Retrospective analysis reveals that incorporating these model uncertainty estimates into the compound selection process could further enhance the model’s hit rates. Lastly, we have experimentally profiled an additional 50 compound-kinase pairs to assess the model’s accuracy in predicting inactive compounds, an often overlooked yet crucial aspect, especially in designing compounds that avoid toxic anti-targets. In this task, the model achieved a negative predictive value of 78%.

## 2 Results

### 2.1 IC50’s are accurately recovered from point-of-concentration measurements

Due to its low cost, percentage inhibition remains the most prevalent POC activity measurement in compound-kinase interaction studies. It is theoretically compatible with the IC_50_ metric which is derived through curve fitting based on percentage inhibition data points at multiple compound concentrations. However, outside the IC_50_ context, percentage inhibition is typically determined at only a few (one to three) compound concentrations. This limitation restricts the applicability of curve fitting methodologies for extracting a more robust metric of compound-kinase activity.

Here, we start by demonstrating our ability to accurately predict pIC_50_ values, i.e., *−* log_10_(IC_50_), using percentage inhibition measurements obtained at just a few points of concentration. To do this, we first create bins of concentrations ranging from *<* 100 nM to *≥* 10000 nM, and select compound-kinase pairs for which at least two percentage inhibition measurements in different bins have been collected. We then construct feature vectors that contain at index *i* the measured percentage inhibition value at bin *i* (if it is measured), and otherwise a special dummy value representing N/A if the value is not available. This is illustrated in the bottom panel of Fig. 1A. To train a POC*→*pIC_50_ model, we further select among these pairs those that additionally have a measured pIC_50_ value. This results in 1563 compound-kinase pair examples, which we further split into training and validation sets of size 1329 and 234, respectively. Our compound-kinase activity dataset used throughout this work was carefully curated based on information from the ChEMBL [MGB^+^19] and PubChem [KCC^+^23] databases, as outlined in Section 3.1. For the POC*→*pIC_50_ prediction task, we train a random forest regression model using the featurized percentage inhibition values to predict the measured pIC_50_ value (refer to Section 3.2). Performance of this model is visualized in Fig. 2A. We obtain a validation set root mean squared error (RMSE) of 0.704, and a Spearman rank correlation between predicted and measured pIC_50_ values of 0.820, suggesting the model is a strong predictor of pIC_50_ .

**Figure 2.**
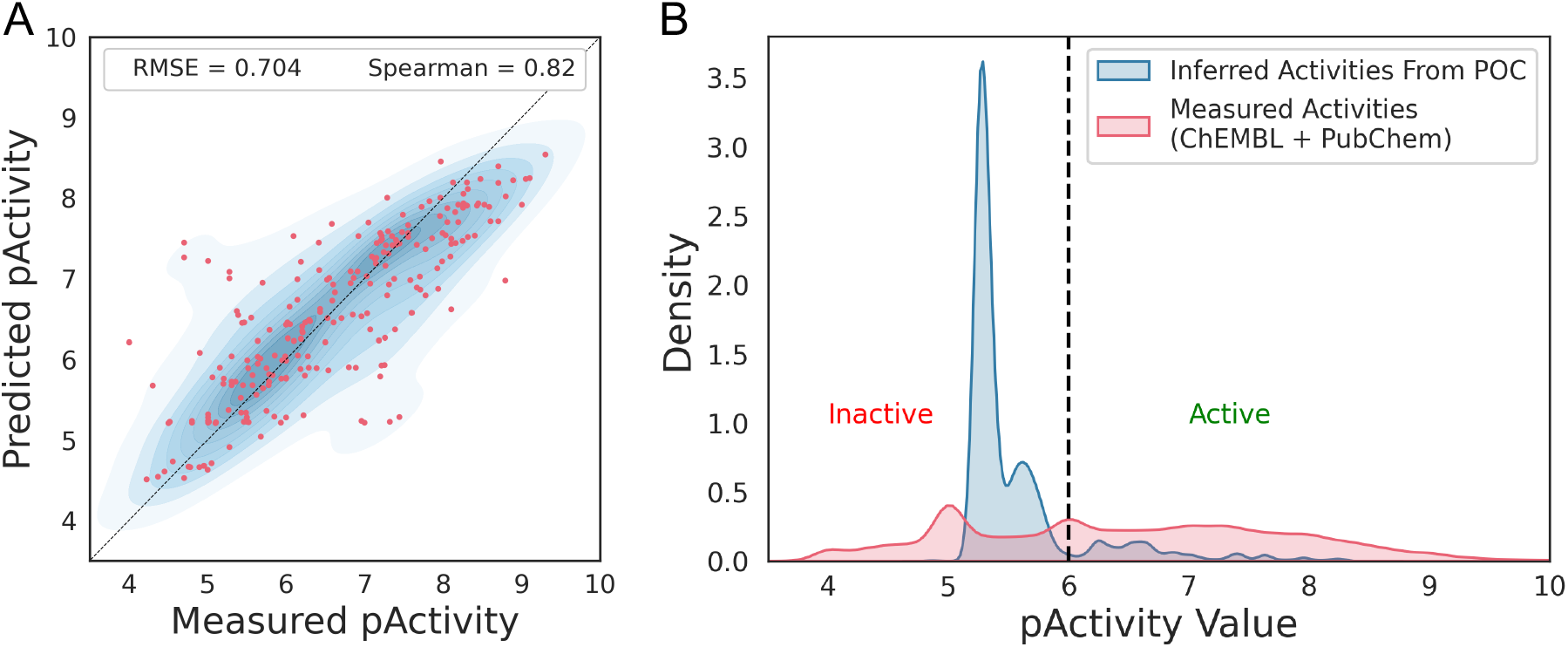
**(A)** Performance of the POC to IC_50_ activity prediction model on the validation set. pIC_50_ values can be accurately predicted from only a few points of concentration. **(B)** Distribution of measured versus inferred activities. We observe that a significantly larger fraction of compound-kinase pairs with inferred activity values are predicted to be inactive, compared to the baseline distribution of activity values. Refer to Supplementary Fig. 2 for the predicted pActivities plotted against the measured percentage inhibition.

Using the trained POC*→*pIC_50_ model, we can now generate inferred pIC_50_ values for compound-kinase pairs that have only a few measured percentage inhibition values (specifically, measurements in at least two different concentration bins), but *no* measured IC_50_, K_i_ or K_d_ . Using our dataset, we are able to extract approximately 70,000 such unlabelled compound-kinase pairs (see Section 3.1). In Fig. 2B, we plot the distribution of predicted pActivity values for this set (shown in blue), along with the distribution of measured pIC_50_ values from ChEMBL and PubChem (in red). Notably, compared to the background distribution, the majority of the inferred activity values are predicted to be inactive (defined here as pActivity *≤* 6, or equivalently, activity *≥* 1000 nM). This augments our dataset with a large number of negative examples indicating a lack of compound-kinase binding. This is not surprising given that compounds found inactive in initial single-dose assays are typically not subjected to further dose-response profiling. However, it’s important to highlight that within this framework, we still identify 13% of the compound-kinase pairs as inferred active (pActivity *>* 6), suggesting potential interactions worth further investigation.

### 2.2 Inferred compound-kinase pairs improve kinome binding predictor performance

We next assess whether integrating inferred pActivity values with the experimentally measured ones enhances the performance of kinome binding predictor models. We benchmarked five models based on varied learning principles, including pairwise kernel ridge regression (pwkrr) [SS02, PAS^+^13], random forest [Bre01], and three deep learning-based methods from the literature: BiMCA [BHS^+^21], DeepDTA [O18], and ConPLex [SSB^+^23]. For the pwkrr, random forest, and ConPLex models, we utilized ECFP4 compound fingerprints. Conversely, the BiMCA and DeepDTA were applied directly to compound SMILES strings. In terms of kinase features, the pwkrr, BiMCA, and DeepDTA were trained on 85-residue binding pocket sequences. On the other hand, the ConPLex model’s kinase features were generated using a pretrained ProtBert protein language model [EHD^+^21]. The random forest model was built separately for each kinase, and thus it does not rely on kinase features. For a more detailed description of all the methods, refer to Section 3.3. We train each model using either only experimentally measured pActivities (for the single-stage baseline model) or a combination of experimentally measured and inferred pActivities (for the two-stage model). When evaluating the model’s performance, we focus on three practical prediction scenarios (see Fig. 1C):

- Predicting the activities of new compound-kinase pairs, where both the individual compound and kinase are present in the training data, but the specific pair under consideration is not. This scenario corresponds to filling in the gaps in an otherwise known compound-kinase interaction matrix (‘ck split’).
- Predicting the activities of new compounds, where the compound itself is not present in the training data, although similar compounds may be included (‘compound split’).
- Predicting the activities of new compound scaffolds, which represents the most challenging scenario. Here, neither the compound in question nor similar compounds within the same cluster are present in the training data (‘scaffold split’).

The results, summarized in Tables 1 and 2, highlight the improvement in predictive performance achieved through the integration of POC data. Our two-stage approach consistently improves performance across the prediction scenarios and learning algorithms evaluated. The only exception is the random forest model under a ‘compound split’ scenario, where the baseline model slightly outperforms the two-stage model, but with the difference in Spearman correlation being marginal, at the third decimal place. Notably, as evidenced by the differences in evaluation metrics and supported by rigorous permutation testing (refer to Section 3.4), the more challenging the prediction scenario, the greater the improvement in performance achieved by the two-stage model compared to the baseline. For example, when predicting the activities of new compound scaffolds, the top-performing pwkrr two-stage model achieved a Spearman correlation of 0.619, compared to 0.593 achieved by the baseline model. The improvement in performance is notable across the chemical space, with the per-compound cluster difference in Spearman correlation due to POC data integration reaching up to 0.4 (Fig. 3A). Visual inspection of clusters highlighted in panel A of Fig. 3 reveals a tight grouping of compounds with common scaffolds, exit vectors, and essentially structural changes within a potential SAR series (Fig. 3B). For example, while 4-amino-7-alkoxyquinazolines are common scaffolds in clusters 84 and 77, they are distant in the t-SNE space, and it is noteworthy that the latter cluster includes ligands with a unique feature, that is, covalent warheads of the acrylamide type. This analysis highlights the improved capability of the two-stage model in identifying detailed variations among kinase inhibitors.

**Table 1:** Validation set Spearman correlation between measured and predicted pActivities for five benchmarked models across three prediction scenarios. P-values comparing two-stage (‘combo’) and single-stage (‘single’) model results were calculated using permutation tests (refer to Section 3.4). Significant p-values are highlighted in green, and metric values for the best-performing model under each prediction scenario are bolded. Higher Spearman correlation values indicate better model performance.

| model | ck split |  |  | compound split |  |  | scaffold split |  |  |
| --- | --- | --- | --- | --- | --- | --- | --- | --- | --- |
|  | combo | single | p-val | combo | single | p-val | combo | single | p-val |
| pwkrr | 0.870 | 0.870 | 0.6574 | 0.837 | 0.837 | 0.5842 | **0.619** | 0.593 | 0.0000 |
| random forest | 0.844 | 0.843 | 0.3526 | 0.845 | **0.846** | 0.9030 | 0.611 | 0.599 | 0.0000 |
| BIMCA | 0.832 | 0.832 | 0.5413 | 0.767 | 0.764 | 0.1201 | 0.451 | 0.439 | 0.0011 |
| DeepDTA | 0.846 | 0.843 | 0.0535 | 0.774 | 0.763 | 0.0001 | 0.487 | 0.433 | 0.0000 |
| ConPLex | **0.880** | 0.877 | 0.0004 | 0.821 | 0.817 | 0.0030 | 0.568 | 0.560 | 0.0011 |

**Table 2:** Validation set RMSE between measured and predicted pActivities for five benchmarked models across three prediction scenarios. P-values comparing two-stage (‘combo’) and single-stage (‘single’) model results were calculated using permutation tests (refer to Section 3.4). Significant p-values are highlighted in green, and metric values for the best-performing model under each prediction scenario are bolded. Lower RMSE values indicate better model performance.

| model | ck split |  |  | compound split |  |  | scaffold split |  |  |
| --- | --- | --- | --- | --- | --- | --- | --- | --- | --- |
|  | combo | single | p-val | combo | single | p-val | combo | single | p-val |
| pwkrr | 0.657 | 0.662 | 0.0047 | 0.708 | 0.710 | 0.1150 | **1.032** | 1.062 | 0.0000 |
| random forest | 0.706 | 0.707 | 0.2779 | 0.691 | **0.686** | 0.9590 | 1.079 | 1.082 | 0.1694 |
| BIMCA | 0.741 | 0.747 | 0.0600 | 0.839 | 0.847 | 0.0218 | 1.241 | 1.251 | 0.0038 |
| DeepDTA | 0.703 | 0.713 | 0.0172 | 0.822 | 0.841 | 0.0000 | 1.222 | 1.267 | 0.0000 |
| ConPLex | **0.638** | 0.647 | 0.0001 | 0.751 | 0.759 | 0.0027 | 1.153 | 1.175 | 0.0000 |

**Figure 3.**
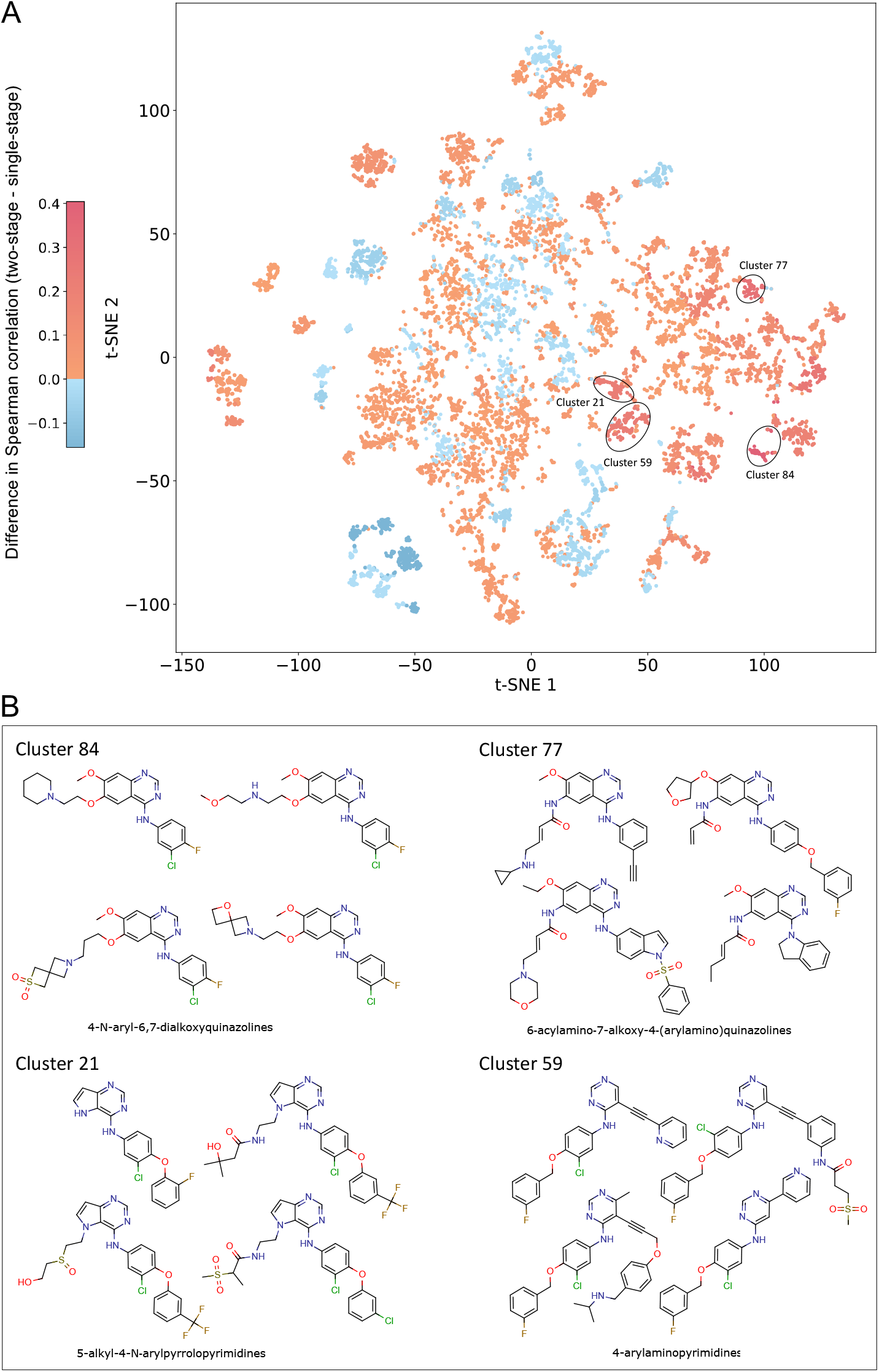
**(A)** Difference in Spearman correlation per compound cluster on the validation set under the ‘scaffold split’ prediction scenario, comparing the two-stage pairwise kernel regression model (pwkrr) with its single-stage counterpart. A higher value indicates superior performance of the two-stage model over the single-stage model. Compounds were first clustered using *k*-means on ECFP4 fingerprints, and t-SNE was applied for visualization (refer to Section 3.4). We highlight selected compound clusters with the largest differences in Spearman correlation, and example structures are shown **(B)**. See Supplementary Fig. 1 for the difference in RMSE.

Interestingly, our results reveal that while a recent deep learning ConPLex method surpasses other models in the easiest scenario of predicting the activities of new compound-kinase pairs, the simpler random forest and kernel learning models substantially outperform all deep learning models benchmarked in the more challenging tasks of predicting the activities of new compounds and compound scaffolds (Tables 1 and 2). It is important to note this finding, as many newly introduced activity prediction methods are currently benchmarked against other published deep learning models, often neglecting comparisons with simpler approaches. To ensure fair comparison across various models, in our experiments, the validation set remained the same across all models within each respective prediction scenario.

Lastly, we assessed the potential for performance improvement in predicting compound selectivity with the two-stage model. To do this, we used the dataset from the Davis *et al*. study [DHH^+^11], which includes dose-response measurements for 72 compounds across approximately 400 kinases. We trained both single-stage and two-stage pwkrr models, excluding all compounds from the Davis *et al*. dataset. ‘True’ selectivity was defined as 1 minus the fraction of kinases each compound binds at *≤* 1000 nM. This metric was correlated against the predicted selectivity (calculated in a similar manner using predicted activities) for both models. The single-stage model achieved a Spearman correlation of 0.542, compared to 0.581 for the multi-stage model. Although this difference is substantial, it is not statistically significant (p-value: 0.2156, permutation test), which could be attributed to the very small number of compounds in the validation set. Nevertheless, the results highlight the multi-stage model’s improved capability in predicting kinase inhibitor selectivity, likely due to the inclusion of POC compound measurements that are often evaluated across multiple kinases, with only a few selected compound-kinase pairs advancing to dose-response testing.

### 2.3 Experimental testing demonstrates practical model utility in early stage drug discovery

We next utilize the top-performing pwkrr two-stage model to screen large purchasable compound libraries against 13 kinases which have a varying number of available training data points (ACVR1, BTK, CSF1R, EGFR, ERBB2, FLT3, IRAK1, IRAK4, JAK2, MERTK, MKNK1, PIK3CA, SYK). For example, EGFR has roughly 6000 data points in the training set, while ACVR1 has only about 200, making it a more challenging target to model. Using a combination of 276 percentage inhibition assays at a compound concentration of 1000 nM (DiscoverX’s KINOMEscan-scanELECT) and 21 K_d_ assays (DiscoverX’s KINOMEscan-KdELECT, see Section 3.5 for the experimental protocol), we experimentally measured 297 previously untested compound-kinase interactions with a predicted pActivity *≥* 6 (or equivalently *≤* 1000 nM), aiming to validate our computational predictions. The Supplementary Data provides the SMILES for compounds, along with UniProt IDs and HGNC symbols for the kinases tested, and includes both the computational predictions and the corresponding results from experimental assays. Supplementary Fig. 3 illustrates the distribution of experimentally measured activity values across all kinases included in the assays, whereas Supplementary Fig. 4 shows the distribution for each kinase separately.

Considering measured K_d_ *≤* 1000 nM and percentage inhibition at 1000 nM *≥* 75%, we attained a hit rate of 40%, which significantly exceeds the average success rates reported in conventional virtual screening endeavors, which often hover between 5% and 25% [ZCS^+^13, LWB^+^19]. Our hit rate, though reduced, remains notably high at 33% when evaluating a more challenging subset of 142 compound-kinase pairs where neither the pair nor the compound overlaps with the training dataset (‘new compounds’). Fig. 4A displays the hit rates as a function of varying percentage inhibition thresholds. Even at the most stringent thresholds, both hit rates remain around 30%. It is worth noting that seven experimentally confirmed novel compound-kinase interactions, spanning seven distinct compounds and five kinases, would have been overlooked by a baseline single-stage model (see Supplementary Data). Among these, four compounds lack very close neighbors in the training dataset with an ECFP4-based Tanimoto similarity to the nearest training compound ranging from 0.55 to 0.73.

**Figure 4.**
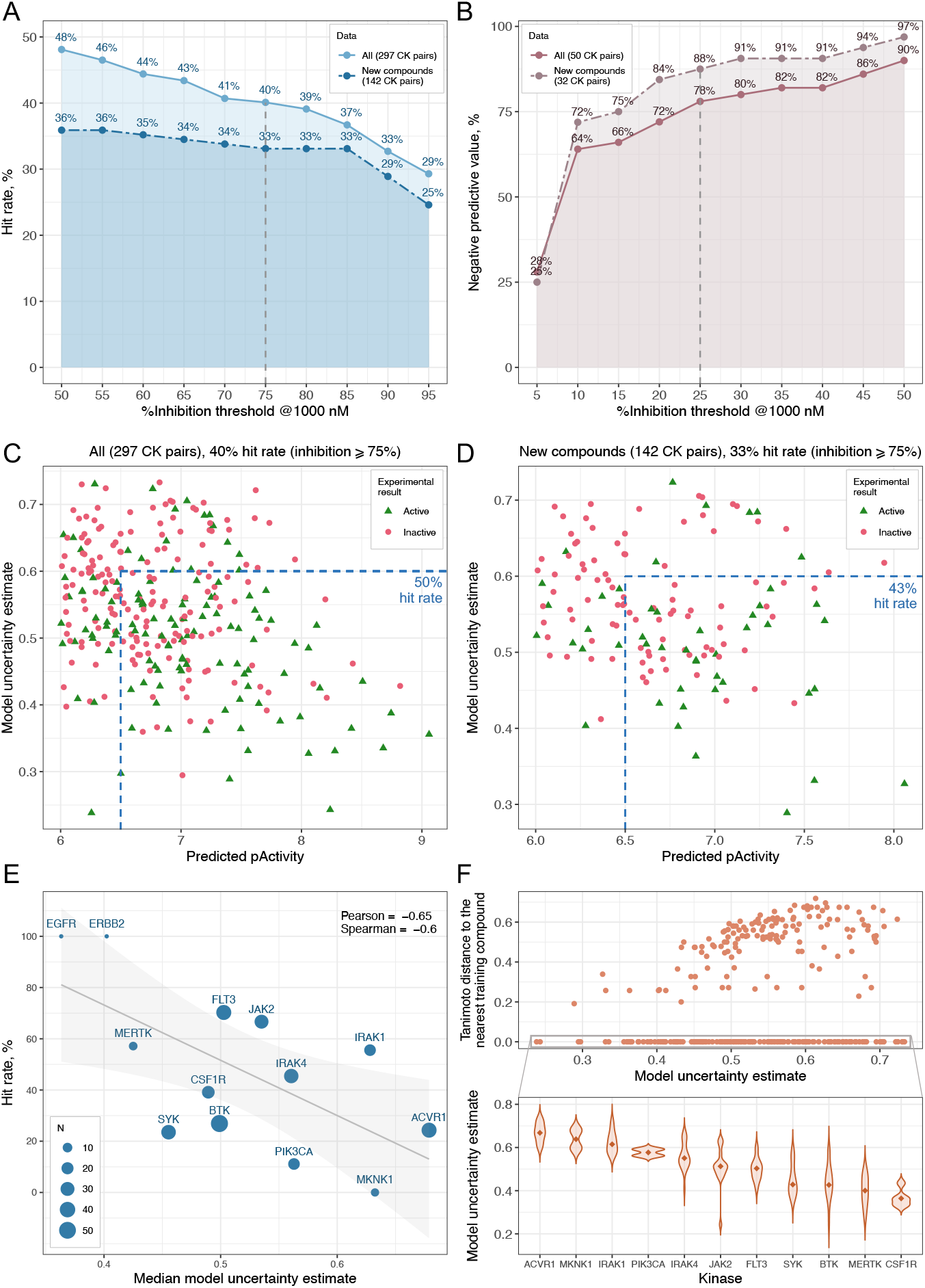
Experimental results: Hit rate **(A)** and negative predictive value **(B)** as functions of the varying percentage inhibition thresholds for all experimentally measured compound-kinase pairs (‘all’), and subsets of pairs where neither the pair nor the compound overlaps with the training dataset (‘new compounds’). The K_d_ threshold for defining actives and inactives remains unchanged (*≤* 1000 nM for actives and > 1000 nM for inactives). **(C)** Two-stage pwkrr model uncertainty estimate plotted versus predicted pActivity for all predicted-as-active compound-kinase pairs and their ‘new compounds’ subset **(D)**. Green triangles indicate validated actives, and red dots denote inactives. The inclusion of model uncertainty estimates in the compound-kinase selection process, as indicated by dashed blue lines, could increase hit rates by 10 percentage points. **(E)** Scatter plot showing the hit rate percentage against the median model uncertainty estimate for each kinase. The size of the circles represents the number of compounds tested per kinase. **(F)** Top panel: Scatter plot of the ECFP4-based Tanimoto distance to the nearest training compound versus the model uncertainty estimate for each compound-kinase pair. Bottom panel: Violin plots displaying the distributions of model uncertainty estimates per kinase for compound-kinase pairs with compounds overlapping with the training data (i.e., with a Tanimoto distance of 0), illustrating that the uncertainty estimates are dependent on both compound and kinase information.

Furthermore, we leverage the connection between kernel ridge regression and Gaussian process [RW06] to calculate the metric of model’s uncertainty for each point estimate of compound-kinase activity (see Section 3.3 for details). The lower the value of this metric, the greater the model’s confidence in its prediction. Uncertainty estimates could be incorporated into the compound selection process. For instance, our retrospective analysis demonstrates that increasing the predicted pActivity threshold to *≥* 6.5, while also applying a threshold of *≤* 0.6 for model uncertainty estimates, would raise hit rates to 50% considering all compound-kinase pairs tested, and to 43% for the ‘new compounds’ subset, which corresponds to a 10 percentage points increase (Fig. 4C-D).

We observe that model uncertainty estimates are strongly correlated with hit rates for each kinase. As expected, ‘dark’ kinases [BML^+^21] and those with limited training data, such as MKNK1 and ACVR1, present greater prediction challenges and exhibit higher uncertainty estimates compared to well-studied kinases like EGFR and FLT3 (Fig. 4E). This pattern holds true for compounds as well; typically, the higher the distance of a tested compound from those in the training set, the higher the associated model uncertainty in predicting its activity. However, Fig. 4F reveals exceptions to this trend. The uncertainty estimate depends on both the compound and the kinase information. Therefore, even compounds overlapping with the training data can have higher associated uncertainty for their activity predictions when paired with less-studied kinases.

Lastly, we conducted experimental profiling (DiscoverX’s KINOMEscan-scanELECT, see Section 3.5 for the experimental protocol) of an additional 50 compound-kinase pairs with predicted pActivity *≤* 5.5 to evaluate the model’s performance in predicting inactive compounds. This aspect is often underemphasized, even though it is critical in the context of designing compounds that strategically avoid toxic anti-targets. Supplementary Fig. 5 shows the distribution of all experimentally measured percentage inhibition values, and Supplementary Fig. 6 shows the distribution for each kinase separately. Defining inactives as compounds with percentage inhibition at 1000 nM *≤* 25%, the model achieved a negative predictive value of 78% (Fig. 4B). This high rate of correctly identifying inactive compounds underscores the model’s potential as a reliable tool, not only for identifying promising kinase inhibitor drug candidates but also for effectively ruling out non-viable or potentially harmful compounds.

## 3 Methods

### 3.1 Data

The bioactivity data used in our experiments was retrieved from two public databases: ChEMBL32 [MGB^+^19] and PubChem [KCC^+^23]. We collected a total of 205,545 IC_50_ measurements from 90,091 compounds tested against 462 different human protein kinases. The compounds were standardized, and those failing our cleaning workflow were filtered out (see Section 3.1.1). In cases where multiple activity measurements were present for a compound-kinase pair, we summarized these into a single activity value by taking the geometric mean of the IC_50_ values. After data cleaning and summarizing, we obtained a final dataset comprising 79,075 compounds measured across 462 kinases, for a total of 141,193 compound-kinase pairs.

Additionally, we collected single-dose activity measurements also from the ChEMBL and PubChem databases. Specifically, we collected examples of compound-kinase pairs for which percentage activity and/or percentage inhibition was measured at at least two separate concentrations. Percentage activity values were converted to percentage inhibition by applying the formula 100 *−* %*activity*. This yielded a total of 69,669 compound-kinase pairs, across 302 kinases.

#### 3.1.1 Compound standardization and cleaning workflow

Compounds were first standardized, following a process similar to the ChEMBL structure curation pipeline [BHF^+^20]. The standardized compounds were then filtered using a set of SMARTS filters, which included, among others, SMARTS for reactive groups, phosphates, sugars, macrocycles, etc. (see [PTS^+^23] for a full list of SMARTS). The compounds were also filtered based on their molecular weight, selecting those with a weight between 250 and 670. Compounds reported with a fluorescence label were stripped down to only their parent compounds. Additionally, a filter was applied to exclude staurosporine- and cholesterol-like compounds, as broad-spectrum tool compounds were not of interest to this study.

#### 3.1.2 Prediction scenarios

Three different training and validation data splits were explored, based on the difficulty of prediction tasks (Fig. 1C). First, we created training and validation sets by randomly splitting compound-kinase pairs (‘ck split’). Next, we split the compounds randomly to ensure distinct compounds in training and validation sets (‘compound split’). In the most challenging scenario, compounds were first clustered, and then some clusters were held out for validation (‘scaffold split’). It should be noted that in the ‘ck split’, a compound present in the training set might also appear in the validation set; however, specific *compound-kinase pairs* from the training set will never appear in the validation set.

For the ‘scaffold split’, we construct training and validation sets as follows. First, we perform *k*-means clustering based on ECFP4 fingerprints of all compounds in the dataset. Then, we continue to add clusters of compounds to the validation set until 10% of the molecules have been designated for validation. The remaining compounds (and all associated compound-kinase pairs) are used for training.

### 3.2 POC data integration

At the data integration step, we train a model to learn a mapping from individual POC measurements to a dose-response IC_50_ value. The inputs to the model are vectors containing percentage inhibition values at different binned concentration values. Specifically, an input is *x* = (*x*_1_, …, *x*_*K*_) where *x*_*j*_ corresponds to a percentage measurement (a scalar between 1 and 100), at concentration bin *j*. The bins are defined in nanomolar (nM) units, with thresholds set at *b*_0_ = 0 nM, *b*_1_ = 100 nM, *b*_2_ = 500 nM, *b*_3_ = 1000 nM, *b*_4_ = 5000 nM, *b*_5_ = 10000 nM, *b*_6_ = 50000 nM, *b*_7_ = 100000 nM, *b*_8_ = 1000000 nM, *b*_9_ = *∞*. A concentration falls into bin *j* if it is between *b*_*j*_ and *b*_*j*+1_. If a given input has no measurement in bin *j*, it is assigned a special value *−*1 representing “no measurement”. During training, we restrict to compound-kinase pairs that have 1) percentage inhibition measurements in at least two separate bins, and 2) an associated IC_50_ value. A schematic representation of our approach can be found in Fig. 1A. Because of the presence of missing values in the data, we choose to use a random forest for the data integration step, which can naturally handle this aspect of the data [Bre01].

### 3.3 Second-stage models

In total, five second-stage models were trained and evaluated on training and validation datasets derived from ‘ck split’, ‘compound split’ and ‘scaffold split’ (Section 3.1.2). Two distinct metrics were reported for each model: Spearman correlation, and root mean squared error (RMSE). We assessed the impact of integrating POC data by training the models with the inclusion of inferred IC_50_ values (two-stage model) and without them (baseline single-stage model).

Note that in this section, the terms ‘compound’ and ‘ligand’ as well as ‘protein’ and ‘kinase’ are used interchangeably.

#### Pairwise kernel ridge regression

We use a pairwise kernel ridge regression model [SS02, PAS^+^13], that operates on an input protein-ligand pair (*p, l*), where the protein *p* is represented as an 85-residue kinase binding pocket sequence retrieved from the KLIFS database [KdGW^+^21], and the ligand *l* is represented as a 1024-bit ECFP4 fingerprint computed using the RDKit library. For a given input (*p, l*), the model’s activity predictions are computed as

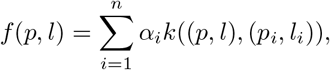

for training protein-ligand pairs (*p*_1_, *l*_1_), …, (*p*_*n*_, *l*_*n*_). The pairwise kernel *k* operating on protein-ligand pairs is defined by the product of a protein kernel and a ligand kernel:

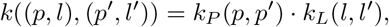

where *k*_*P*_ is calculated based on the (normalized) Striped-Smith-Waterman sequence alignment, and *k*_*L*_ is the Tanimoto kernel. Note that as both *k*_*P*_ and *k*_*L*_ are bounded between 0 and 1, the pairwise kernel *k* is also bounded within this range. The parameters *α*_1_, …, *α*_*n*_ are fit by minimizing a standard kernel ridge regression objective using the conjugate gradient method.

The pairwise kernel ridge regression model also admits a convenient interpretation as a Gaussian process associated with the kernel *k* [RW06]. This means that we can naturally compute an uncertainty estimate associated with each activity prediction. Specifically, since our pairwise kernel is 1 for identical compound-kinase pairs, the expression for the variance of a new compound-kinase pair (*p, l*) is given by

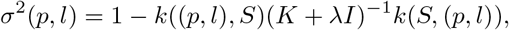

where *K* is the *n × n* training kernel, and *k*(*S*, (*p, l*)) is the *n*-dimensional vector whose *i*th entry is *k*((*p*_*i*_, *l*_*i*_), (*p, l*)) for the *i*th training example (*p*_*i*_, *l*_*i*_).

#### Random forest

We use a standard random forest model as implemented in scikit-learn software package [PVG^+^11]. Rather than designing a single model that takes protein-ligand pairs as inputs, we fit separate random forests for each kinase. For ligands, we use 1024-bit ECFP4 fingerprints computed using the RDKit library.

#### DeepDTA

The DeepDTA [O18] architecture consists of two embedding modules: one ligand embedding module, and a second protein embedding module. Both embedding modules share the same architecture, consisting of a series of convolutional layers, followed by a pooling layer to obtain sequence-level embeddings from compound SMILES and 85-residue kinase binding pocket sequence strings, respectively. After embedding a SMILES string and an amino acid sequence, the resulting feature vectors are concatenated, and passed through a series of linear layers interspersed with dropout layers to obtain the final scalar output.

#### BiMCA

Similar to DeepDTA, BiMCA [BHS^+^21] uses convolutional neural network layers to learn feature embeddings from both compound SMILES and protein sequence strings. Then, unlike DeepDTA, BiMCA uses context attention layers to fuse information from both modalities, allowing the ligand representation access to contextual information from the protein embedding, and vice versa. Finally, the resulting feature vectors are concatenated and passed through a fully connected module to produce a scalar output.

#### ConPLex

ConPLex [SSB^+^23] is another recent neural network-based model used to predict compound-kinase binding affinity. ConPLex featurizes ligands using ECFP4 fingerprints, and kinases using a pre-trained ProtBERT language model [EHD^+^21]. The model then uses fully connected layers with ReLU activations to project compound and kinase features into a shared embedding space. From this shared space, binding affinity between a compound and a kinase is estimated by computing a dot product between the compound and kinase embeddings. Unlike the other models considered here, ConPLex also employs contrastive learning stage, wherein the model is trained to simultaneously predict bioactivities, while maximizing the distance between real drugs and synthetically-generated decoys in embedding space. Here, we used the same features as those reported in their model for binding affinity prediction, along with the same contrastive learning procedure.

### 3.4 Post-analysis on predictions

#### Significance testing

To test the statistical significance of the improvement from data integration, we use a non-parametric permutation test. For every example in the validation set, we make predictions using both the single- and two-stage models, and compute the difference in performance metrics between the two models. Then, to generate a null distribution, we randomly permute single- and two-stage labels 10,000 times across examples in the validation set, and calculate the difference in each performance metric for each permutation. The observed differences are then compared against the null distribution to calculate p-values for each metric and model.

#### Compound-wise analysis

To further analyze the impact of integrating inferred IC_50_ values on predicting the activities of structurally diverse compounds, we applied *k*-means clustering to the compounds in the validation set for the ‘scaffold split’ scenario, using ECFP4 fingerprints. We set the number of clusters at 100 to capture potential common scaffolds within each cluster. After clustering, we calculated the differences in performance metrics (Spearman correlation and RMSE) between the single-stage and two-stage models for compound-kinase pairs associated with compounds in each cluster. Additionally, for visualization purposes, we first applied PCA to the compound ECFP4 fingerprints, and then we employed t-SNE on the 20 principal components (Fig. 3A and Supplementary Fig. 1).

### 3.5 Experimental profiling

We experimentally profiled a total of 347 compound-kinase pairs, encompassing 13 kinases (ACVR1, BTK, CSF1R, EGFR, ERBB2, FLT3, IRAK1, IRAK4, JAK2, MERTK, MKNK1, PIK3CA, SYK) and 139 compounds (see Supplementary Data for a list of compound SMILES), based on predictions from the top-performing two-stage pwkrr model (see Section 2.3). To generate new dissociation constant (K_d_) and percentage inhibition values, we sent the compounds to DiscoverX (Eurofins Corporation) for KINOMEscan profiling service. The KINOMEscan screening platform utilizes an active site-directed competition binding assay to measure interactions between test compounds and selected human kinases, without the need for ATP. This technique hinges on the principle that compounds binding to the kinase active site prevent the kinase’s interaction with the immobilized active-site directed ligand, and therefore result in a diminished amount of kinase captured on the solid support [FBIT^+^05].

In our experiments, K_d_ determination was conducted using the KdELECT method (https://www.eurofinsdiscovery.com/solution/kdelect), while percentage inhibition at a compound concentration of 1000 nM, relevant in the context of kinase inhibition, was measured using the scan-ELECT protocol (https://www.eurofinsdiscovery.com/solution/scanelect). Both methods are parts of the KINOMEscan platform.

#### 3.5.1 Protocol description

Kinase-tagged T7 phage strains were grown in an *E. coli* host derived from the BL21 strain. The *E. coli* were grown to log-phase, infected with T7 phage (multiplicity of infection = 0.4), and incubated with shaking at 32*?*C until lysis occurred (90-150 minutes). The lysates were then centrifuged (6000 x g) and filtered to remove cell debris. The remaining kinases were produced in HEK-293 cells and subsequently tagged with DNA for qPCR detection.

Streptavidin-coated magnetic beads were treated with biotinylated small molecule ligands for 30 minutes at room temperature to generate affinity resins for kinase assays. In order to remove unbound ligand and reduce non-specific binding, the ligand-bound beads were then blocked with excess biotin and washed with a blocking buffer (SeaBlock (Pierce), 1% BSA, 0.05% Tween 20, 1 mM DTT). Binding reactions were constructed by mixing kinases, ligand-bound beads, and test compounds in 1x binding buffer (20% SeaBlock, 0.17x PBS, 0.05% Tween 20, 6 mM DTT).

Test compounds for percentage inhibition assays were prepared as 40x stocks in 100% DMSO, whereas for K_d_ assays, they were prepared as 111x stocks in 100% DMSO. K_d_ ‘s were determined using an 11-point 3-fold compound dilution series with three DMSO control points. Prepared compounds were directly diluted into the assays. All reactions were carried out in polypropylene 384-well plates in a final volume of 0.02 ml. Following incubation at room temperature with shaking for one hour, the affinity beads were washed with a wash buffer (1x PBS, 0.05% Tween 20). Subsequently, the beads were resuspended in an elution buffer (1x PBS, 0.05% Tween 20, 0.5 *µ*M non-biotinylated affinity ligand), and incubated at room temperature with shaking for 30 minutes. Finally, the concentration of each kinase in the eluates was measured using qPCR.

#### Determination of percentage inhibition and K_d_

In case of percentage inhibition assays, test compounds were screened at a single concentration of 1000 nM, and the percentage inhibition of a kinase was calculated as follows:

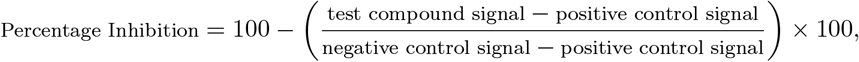

where the negative control is DMSO (0% inhibition) and the positive control is the control compound (100% inhibition).

K_d_ ‘s were calculated with a standard dose-response curve using the Hill equation:

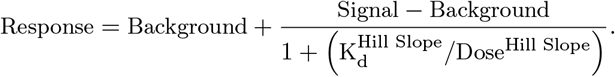

The Hill Slope was set to -1, and curves were fitted using a non-linear least square fit with the Levenberg-Marquardt algorithm [Hil10, Lev44].

## 4 Discussion

The heterogeneity in available compound-kinase activity data calls for machine learning approaches capable of integrating various experimental readouts during model training. Here, we have introduced a two-stage machine learning framework, which, to the best of our knowledge, is the first to leverage the typically overlooked POC data alongside dose-response activities for compound-kinase binding prediction. We have demonstrated that our approach is adaptable to various learning algorithms and molecular descriptors. The improvement in performance is evident across all the algorithms evaluated here, especially in the most challenging and practical early-stage drug discovery tasks of predicting activities of new compounds and compound scaffolds that were not seen in the training data. This represents an advancement in navigating the complex compound-kinase interaction space, facilitating more effective exploration of a broader chemical spectrum. Due to the integration of POC screening results, often carried out across larger kinase panels, our two-stage approach also enhances the prediction of kinase inhibitor selectivity.

As experimental testing is the ultimate method for assessing a model’s utility in drug discovery efforts, we have profiled a total of 347 compound-kinase pairs based on activity predictions from the top-performing two-stage pairwise kernel regression model, achieving a hit rate of 40% and a negative predictive value of 78%. Our hit rate is notably higher than those typically reported in virtual screening studies, which often range between 5% and 25% [ZCS^+^13, LWB^+^19]. Our experimentally-generated data is available to the community alongside this publication. We also derived uncertainty estimates associated with kernel model activity predictions and demonstrated how they could guide the compound selection process, leading to improved hit rates.

Our study underscores the significance of thoroughly assessing the applicability domain of activity prediction machine learning models. An understanding of each model’s capacity to generalize to previously unseen data is of utmost importance, and can be achieved by meticulously constructing training and validation splits with careful consideration of compound and kinase overlaps between them [BSS^+^22]. This is especially important given the intended application of the model. For instance, in our study, the ConPLex method exhibited superior performance in filling gaps in tested compound-kinase interaction matrices. This is relevant, for example, in preparing activity data for other downstream tasks where missing values are not allowed. Conversely, in screening new compound libraries, random forest and kernel learning outperformed all deep learning approaches evaluated here, demonstrating a notable Spearman correlation difference on the same validation set of up to 0.17. This shows that despite the increasing popularity of deep learning algorithms in recent years, it is crucial to compare these advanced methods not only against their counterparts but also against simpler, yet still powerful, more traditional models. This would ensure a comprehensive evaluation, highlighting the strengths and limitations of each method in various contexts. Randomly splitting compound-kinase pairs into training and validation sets results in overoptimistic performance in terms of generalization to previously unseen data, as was also observed in other work [OKK23].

Furthermore, constructing high-quality training data is essential. In the literature, models are frequently trained and evaluated on kinase datasets such as those from Anastassiadis *et al*. [ADD^+^11], Davis *et al*. [DHH^+^11], and Metz *et al*. [MJS^+^11] studies. While these datasets are of high quality, they represent a limited chemical spaces, typically including only up to a few hundred compounds. Here, we have curated a kinome-wide dataset from ChEMBL and PubChem, comprising roughly 80,000 compounds that have undergone rigorous cleaning workflow. This process standardizes compounds and filters out structures with undesirable characteristics. These include, among others, reactivity, staurosporine-like structures, the presence of long aliphatic chains, or atoms such as Si, Se, and I, as well as fragments prone to rapid oxidation (see Section 3.1.1). We advocate for meticulous data preparation to ensure reliable and effective machine learning applications in kinase research.

In this study, for simplicity, we used only IC_50_ values from the available dose-response data during model training. However, our methodology is equally capable of incorporating other commonly used readouts, such as K_d_ and K_i_ values. IC_50_, K_d_, and K_i_ readouts are frequently used together in training models for kinase inhibitor activity prediction. Incorporating these additional activity data could further enhance both the robustness and accuracy of the predictive model. However, further work is required to account for inconsistencies between various assays [LR24]. Even though we focused on kinase targets, we anticipate that our two-stage framework could be applicable to other protein classes with a similar spectrum of data types.

Lastly, while our hit rate significantly exceeds those commonly observed in virtual screening efforts [ZCS^+^13, LWB^+^19], it is important to acknowledge that hit rates will vary based on the specific targets and chemical libraries utilized. Although we primarily used percentage inhibition assays to validate our computational predictions - due to their standard application in initial compound screening for their high throughput and cost-effectiveness - they have their limitations. These assays provide only a snapshot of compound activity at a single concentration under a specific set of experimental conditions, potentially overlooking the complex dynamics and variety of compound-kinase interaction mechanisms. This limitation may result in a partial or misleading evaluation of a compound’s efficacy and selectivity. While single-dose screening is a well-established strategy to identify hits, confirmatory evaluation in dose-response assays is required for accurate compound binding assessment. Additional biochemical profiling experiments can reveal further insight into inhibition mechanisms (e.g., competitive, allosteric or time-dependent inhibition).

In conclusion, we believe that this works emphasizes the importance of careful model evaluation under various prediction scenarios as well as sheds light on the untapped potential of POC experimental readouts in the compound activity modelling tasks. Our approach provides valuable insights into the compound-kinase interaction landscape, and we anticipate that it will enable more efficient and economical development of activity datasets for kinase drug discovery.

## Acknowledgments

The authors would like to thank Marcel Patek for his review of the chemical series within the clustering results, as well as Andreas Bender, Tero Aittokallio, Olivier Elemento, Francesco Trozzi, and Ravi Rajagopalan for providing valuable feedback on the manuscript.

## References

[ADD+11]T. Anastassiadis, S. W. Deacon, K. Devarajan, H. Ma, and J. R. Peterson. Comprehensive assay of kinase catalytic activity reveals features of kinase inhibitor selectivity. Nat. Biotechnol., 29(11):1039–1045, 2011.

[BHF+20]A. P. Bento, A. Hersey, E. Félix, G. Landrum, A. Gaulton, F. Atkinson, L. J. Bellis, M. De Veij, and A. R. Leach. An open source chemical structure curation pipeline using RDKit. J. Cheminform., 12(51):1–16, 2020.

[BHS+21]J. Born, T. Huynh, A. Stroobants, W. D. Cornell, and M. Manica. Active site sequence representations of human kinases outperform full sequence representations for affinity prediction and inhibitor generation: 3D effects in a 1D model. J. Chem. Inf. Model., 62(2):240–257, 2021.

[BML+21]M. E. Berginski, N. Moret, C. Liu, D. Goldfarb, P. K. Sorger, and S. M. Gomez. The Dark Kinase Knowledgebase: an online compendium of knowledge and experimental results of understudied kinases. Nucleic Acids Res., 49(D1):D529–D535, 2021.

[Bre01]L. Breiman. Random forests. Machine Learning, 45:5–32, 2001.

[BSS+22]A. Bender, N. Schneider, M. Segler, W. Patrick Walters, O. Engkvist, and T. Rodrigues. Evaluation guidelines for machine learning tools in the chemical sciences. Nat. Rev. Chem., 6(6):428–442, 2022.

[CCAS+15]I. Cortés-Ciriano, Q. U. Ain, V. Subramanian, E. B. Lenselink, O. Méndez-Lucio, A. P. IJzerman, G. Wohlfahrt, P. Prusis, T. E. Malliavin, G. J. van Westen, and A. Bender. Polypharmacology modelling using proteochemometrics (PCM): recent methodological developments, applications to target families, and future prospects. Med. Chem. Commun., 6(1):24–50, 2015.

[CPS+18]A. Cichonska, T. Pahikkala, S. Szedmak, H. Julkunen, A. Airola, M. Heinonen, T. Aittokallio, and J. Rousu. Learning with multiple pairwise kernels for drug bioactivity prediction. Bioinformatics, 34(13):i509–i518, 2018.

[CRA+21]A. Cichońska, B. Ravikumar, R. J. Allaway, F. Wan, S. Park, O. Isayev, S. Li, M. Mason, A. Lamb, Z. Tanoli, M. Jeon, and et al. Crowdsourced mapping of unexplored target space of kinase inhibitors. Nat. Commun., 12(1):3307, 2021.

[CRP+17]A. Cichonska, B. Ravikumar, E. Parri, S. Timonen, T. Pahikkala, A. Airola, K. Wennerberg, J. Rousu, and T. Aittokallio. Computational-experimental approach to drugtarget interaction mapping: a case study on kinase inhibitors. PLoS Comput. Biol., 13(8):e1005678, 2017.

[DHH+11]M. I. Davis, J. P. Hunt, S. Herrgard, P. Ciceri, L. M. Wodicka, G. Pallares, M. Hocker, D. K. Treiber, and P. P. Zarrinkar. Comprehensive analysis of kinase inhibitor selectivity. Nat. Biotechnol., 29(11):1046–1051, 2011.

[DQJ+22]B. X. Du, Y. Qin, Y. F. Jiang, Y. Xu, S. M. Yiu, H. Yu, and J. Y. Shi. Compound–protein interaction prediction by deep learning: databases, descriptors and models. Drug Discov. Today, 27(5):1350–1366, 2022.

[DSSGP22]G. De Simone, D. S. Sardina, M. R. Gulotta, and U. Perricone. KUALA: a machine learning-driven framework for kinase inhibitors repositioning. Sci. Rep., 12(1):17877, 2022.

[DTME20]L. David, A. Thakkar, R. Mercado, and O. Engkvist. Molecular representations in AI-driven drug discovery: a review and practical guide. J. Cheminform., 12(1):1–22, 2020.

[EHD+21]A. Elnaggar, M. Heinzinger, C. Dallago, G. Rehawi, Y. Wang, L. Jones, T. Gibbs, T. Feher, C. Angerer, M. Steinegger, D. Bhowmik, and B. Rost. Prottrans: Toward understanding the language of life through self-supervised learning. IEEE Trans. Pattern Anal. Mach. Intell., 44(10):7112–7127, 2021.

[FBIT+05]M. A. Fabian, W. H. Biggs III, D. K. Treiber, C. E. Atteridge, M. D. Azimioara, M. G. Benedetti, T. A. Carter, P. Ciceri, P. T. Edeen, M. Floyd, J. M. Ford, and et al. A small molecule–kinase interaction map for clinical kinase inhibitors. Nat. Biotechnol., 23:329–336, 2005.

[Hil10]A. V. Hill. The possible effects of the aggregation of the molecules of hemoglobin on its dissociation curves. J. Physiol., 40:iv–vii, 1910.

[KCC+23]S. Kim, J. Chen, T. Cheng, A. Gindulyte, J. He, S. He, Q. Li, B. A. Shoemaker, P. A. Thiessen, B. Yu, L. Zaslavsky, J. Zhang, and E. E. Bolton. PubChem 2023 update. Nucleic Acids Res., 51(D1):D1373—-D1380, 2023.

[KdGW+21]G. K. Kanev, C. de Graaf, B. A. Westerman, I. J. de Esch, and A. J. Kooistra. KLIFS: an overhaul after the first 5 years of supporting kinase research. Nucleic Acids Res., 49(D1):D562–D569, 2021.

[KZEK23]M. Kalemati, M. Zamani Emani, and S. Koohi. BiComp-DTA: Drug-target binding affinity prediction through complementary biological-related and compression-based featurization approach. PLoS Comput. Biol., 19(3):e1011036, 2023.

[KZK+23]G. K. Kanev, Y. Zhang, A. J. Kooistra, A. Bender, R. Leurs, D. Bailey, T. Würdinger, C. de Graaf, I. J. de Esch, and B. A. Westerman. Predicting the target landscape of kinase inhibitors using 3D convolutional neural networks. PLoS Comput. Biol., 19(9):e1011301, 2023.

[Lev44]K. Levenberg. A method for the solution of certain non-linear problems in least squares. Q. Appl. Math., 2:164–168, 1944.

[LKN+23]C. Liu, P. Kutchukian, N. D. Nguyen, M. AlQuraishi, and P. K. Sorger. A hybrid structure-based machine learning approach for predicting kinase inhibition by small molecules. J. Chem. Inf. Model., 63(17):5457–5472, 2023.

[LLP23]Y. Luo, Y. Liu, and J. Peng. Calibrated geometric deep learning improves kinase–drug binding predictions. Nat. Mach. Intell., 5:1390–1401, 2023.

[LR24]G. A. Landrum and S. Riniker. Combining IC50 or Ki values from different sources is a source of significant noise. J. Chem. Inf. Model., 2024.

[LTZ+23]S. Li, T. Tian, Z. Zhang, Z. Zou, D. Zhao, and J. Zeng. PocketAnchor: Learning structure-based pocket representations for protein-ligand interaction prediction. Cell Syst., 14(8):692–705, 2023.

[LWB+19]J. Lyu, S. Wang, T. E. Balius, I. Singh, A. Levit, Y. S. Moroz, M. J. O’Meara, T. Che, E. Algaa, K. Tolmachova, A. A. Tolmachev, B. K. Shoichet, B. L. Roth, and J. J. Irwin. Ultra-large library docking for discovering new chemotypes. Nat., 566(7743):224–229, 2019.

[MGB+19]D. Mendez, A. Gaulton, A. P. Bento, J. Chambers, M. De Veij, E. Félix, M. P. Magariños, J. F. Mosquera, P. Mutowo, M. Nowotka, M. Gordillo-Marañón, F. Hunter, L. Junco, G. Mugumbate, M. Rodriguez-Lopez, F. Atkinson, N. Bosc, C. J. Radoux, A. Segura-Cabrera, A. Hersey, and A. R. Leach. ChEMBL: towards direct deposition of bioassay data. Nucleic Acids Res., 47(D1):D930–D940, 2019.

[MJS+11]J. T. Metz, E. F. Johnson, N. B. Soni, P. J. Merta, L. Kifle, and P. J. Hajduk. Navigating the kinome. Nat. Chem. Biol., 7(4):200–202, 2011.

[MM12]E. Martin and P. Mukherjee. Kinase-kernel models: accurate in silico screening of 4 million compounds across the entire human kinome. J. Chem. Inf. Model., 52(1):156–170, 2012.

[NPC16]A. C. Nascimento, R. B. Prudêncio, and I. G. Costa. A multiple kernel learning algorithm for drug-target interaction prediction. BMC Bioinformatics, 17:1–16, 2016.

[OKK23]W. J. G. Ong, P. Kirubakaran, and J. Karanicolas. Poor generalization by current deep learning models for predicting binding affinities of kinase inhibitors, 2023. Preprint at https://www.biorxiv.org/content/10.1101/2023.09.04.556234v1.

[PAS+13]T. Pahikkala, A. Airola, M. Stock, B. De Baets, and W. Waegeman. Efficient regularized least-squares algorithms for conditional ranking on relational data. Machine Learning, 93:321–356, 2013.

[PHL+23]H. Park, S. Hong, M. Lee, S. Kang, R. Brahma, K. H. Cho, and J. M. Shin. AiKPro: deep learning model for kinome-wide bioactivity profiling using structure-based sequence alignments and molecular 3D conformer ensemble descriptors. Sci. Rep., 13(1):10268, 2023.

[PTS+23]R. Park, R. Theisen, N. Sahni, M. Patek, A. Cichońska, and R. Rahman. Preference optimization for molecular language models, 2023. arXiv preprint arXiv:2310.12304.

[PVG+11]F. Pedregosa, G. Varoquaux, A. Gramfort, V. Michel, B. Thirion, O. Grisel, M. Blondel, P. Prettenhofer, R. Weiss, V. Dubourg, J. Vanderplas, A. Passos, D. Cournapeau, M. Brucher, M. Perrot, and E. Duchesnay. Scikit-learn: machine learning in Python. J. Mach. Learn. Res., 12:2825–2830, 2011.

[RW06]C. E. Rasmussen and C. K. Williams. Gaussian processes for machine learning. MIT Press, Cambridge, 2006.

[SS02]B. Schölkopf and A. J. Smola. Learning with kernels: support vector machines, regularization, optimization, and beyond. MIT Press, Cambridge, 2002.

[SSB+23]R. Singh, S. Sledzieski, B. Bryson, L. Cowen, and B. Berger. Contrastive learning in protein language space predicts interactions between drugs and protein targets. Proc. Natl. Acad. Sci., 120(24):e2220778120, 2023.

[TAA+22]M. A. Thafar, M. Alshahrani, S. Albaradei, T. Gojobori, M. Essack, and X. Gao. Affinity2Vec: drug-target binding affinity prediction through representation learning, graph mining, and machine learning. Sci. Rep., 12(1):4751, 2022.

[ZCS+13]T. Zhu, S. Cao, P. C. Su, R. Patel, D. Shah, H. B. Chokshi, R. Szukala, M. E. Johnson, and K. E. Hevener. Hit identification and optimization in virtual screening: Practical recommendations based on a critical literature analysis. J. Med. Chem., 56(17):6560–6572, 2013.

[O18]H. Öztürk, A. Özgür, and E. Ozkirimli. DeepDTA: deep drug–target binding affinity prediction. Bioinformatics, 34(17):i821–i829, 2018.

