## Supplementary Information for "Leveraging multiple data types for improved compound-kinase bioactivity prediction"

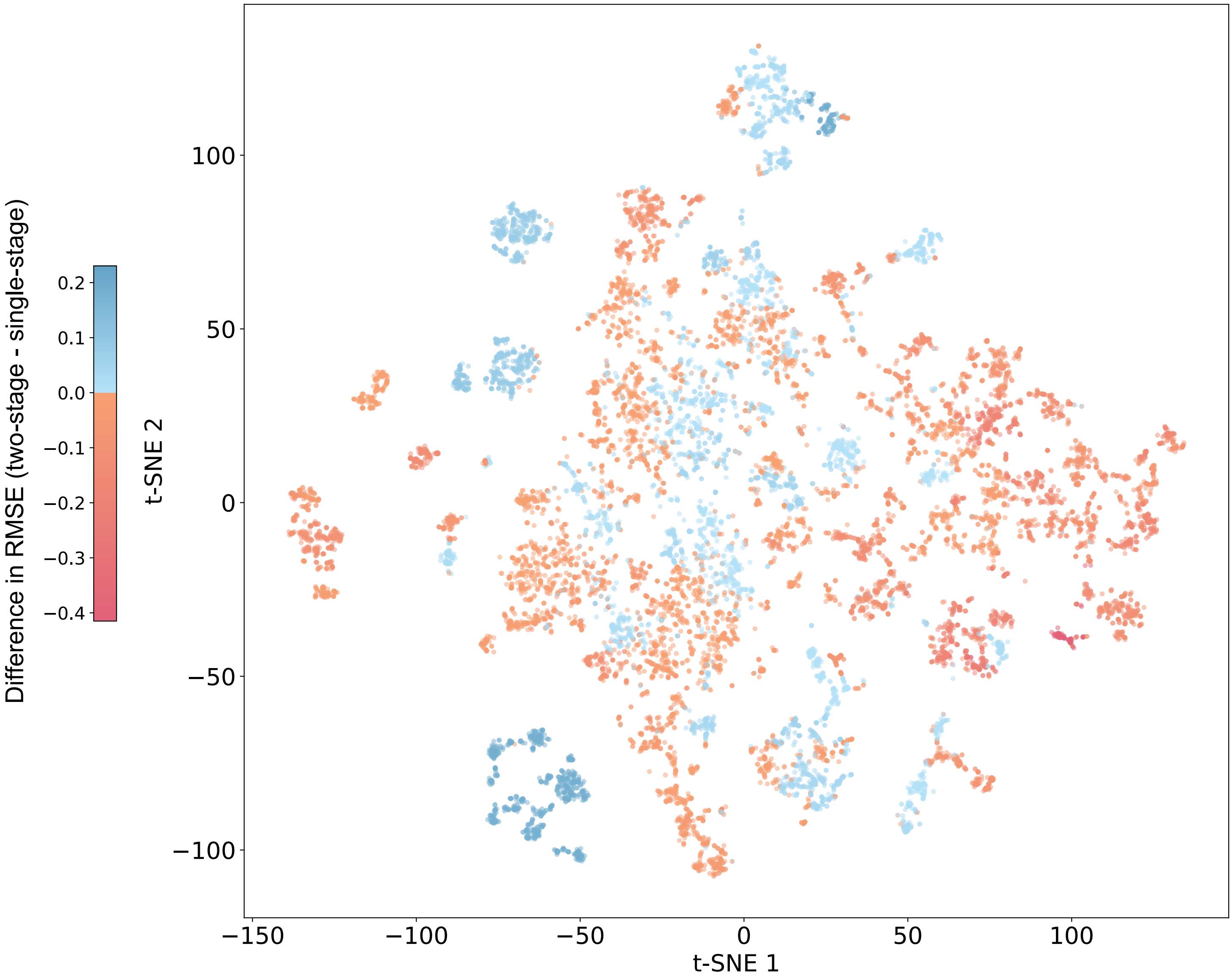


**Supplementary Figure 1**. Difference in RMSE per compound cluster on the validation set under the 'scaffold split' prediction scenario, comparing the two-stage pairwise kernel regression model (pwkrr) with its single-stage counterpart. A lower value indicates superior performance of the two-stage model over the single-stage model. Compounds were first clustered using k-means on ECFP4 fingerprints, and t-SNE was applied for visualization (refer to Section 3.4 of the main publication).

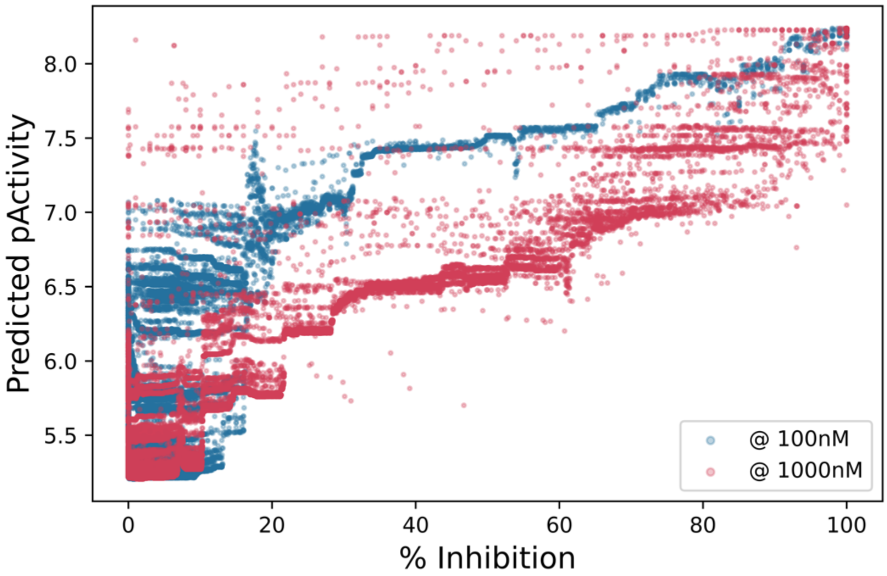


**Supplementary Figure 2**. Predicted pActivity as a function of the measured percentage inhibition at compound concentrations of 100 nM (blue) and 1000 nM (red) across approximately 70,000 compound-kinase pairs with no measured $IC_{50}$, $K_i$, or $K_d$.

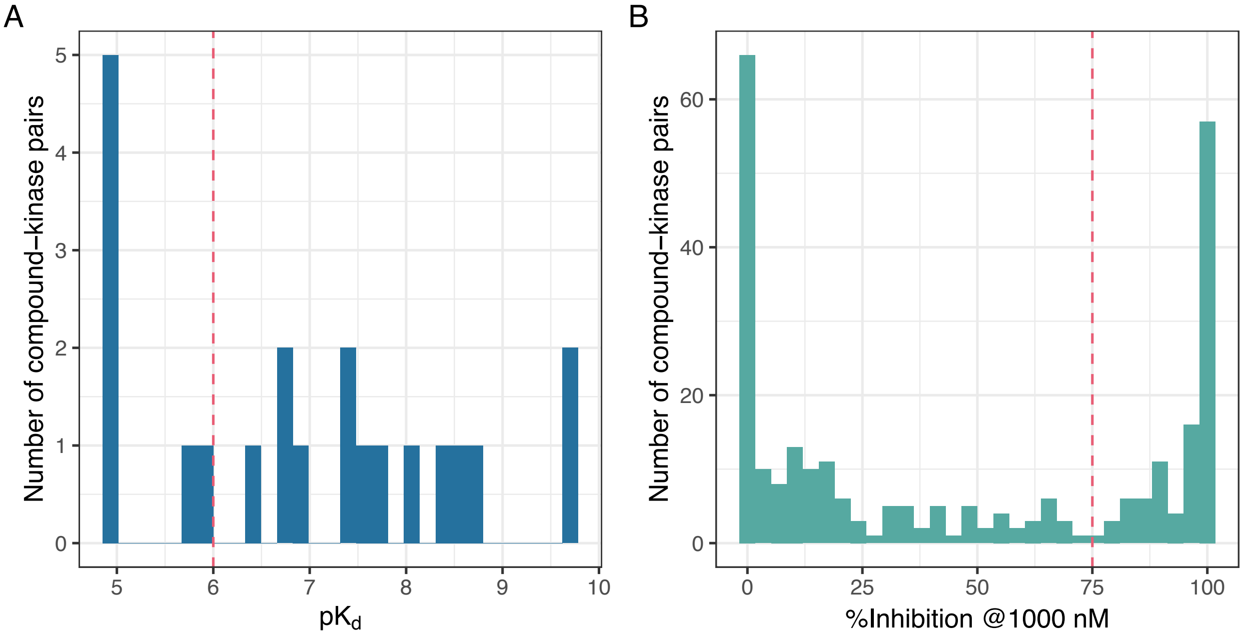


**Supplementary Figure 3**. Distribution of experimentally measured $pK_d$ (**A**) and percentage inhibition at 1000 nM (**B**) across 297 previously untested compound-kinase pairs predicted as active by the two-stage pwkrr model. Red vertical dashed lines serve as references for an activity threshold of 1000 nM ($pK_d$ of 6, in **A**) and 75% inhibition (in **B**).

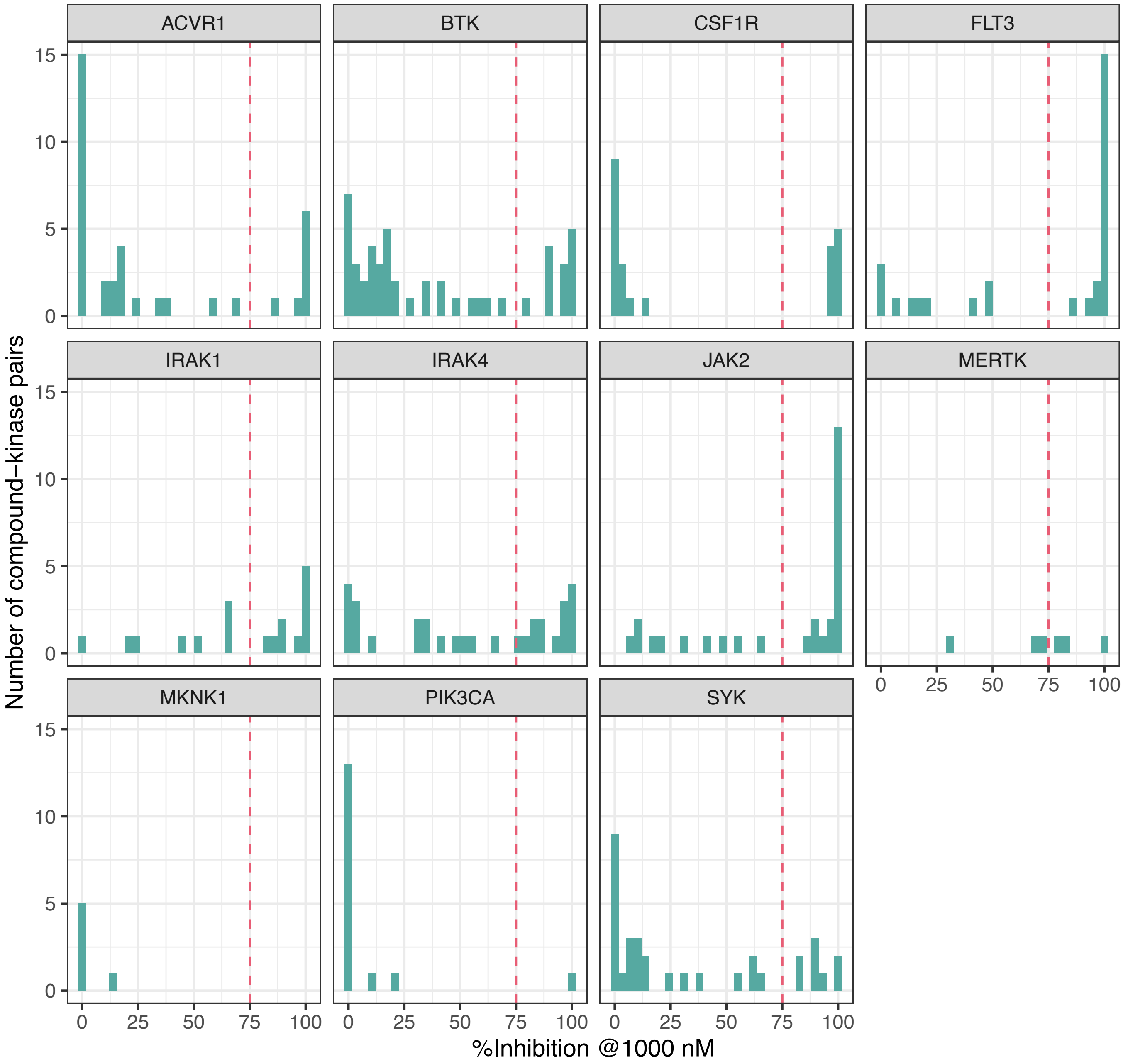


**Supplementary Figure 4**. Distribution of experimentally measured percentage inhibition at 1000 nM per kinase across 276 compound-kinase pairs predicted to be active by the two-stage pwkrr model and tested in single-dose assays. Red vertical dashed lines serve as references for an activity threshold of 75% inhibition.

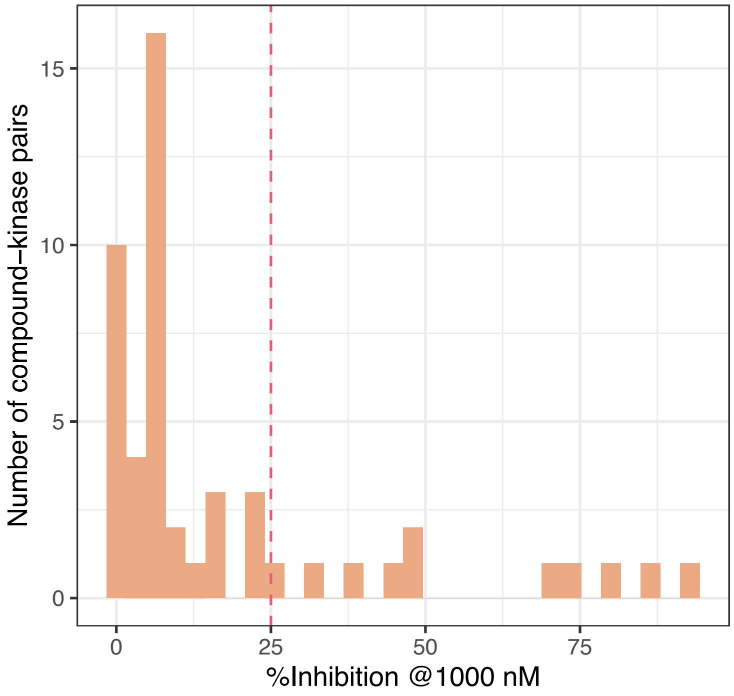


**Supplementary Figure 5**. Distribution of experimentally measured percentage inhibition at 1000 nM across 50 compound-kinase pairs predicted to be inactive by the two-stage pwkrr model. A red vertical dashed line serves as a reference for the inactive threshold of 25% inhibition.

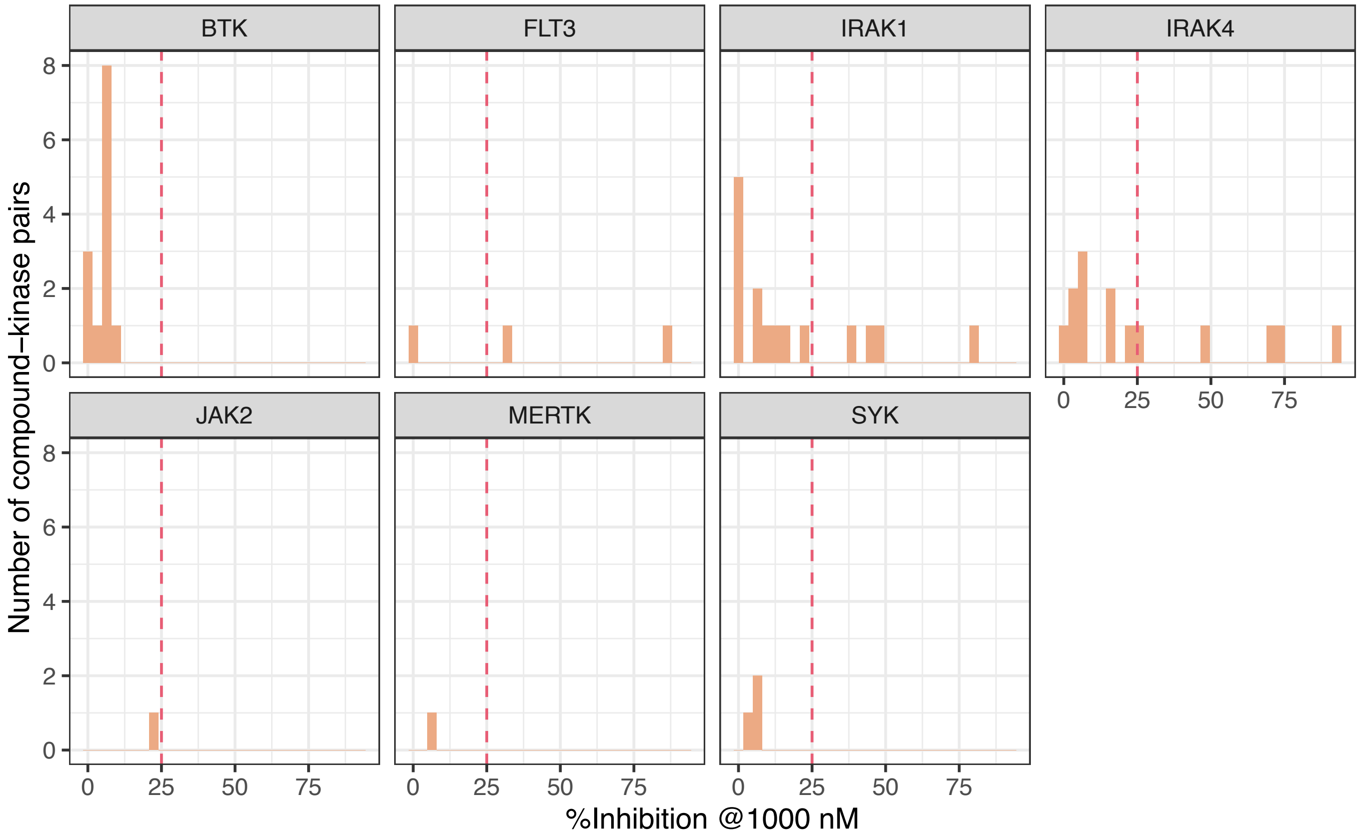


**Supplementary Figure 6**. Distribution of experimentally measured percentage inhibition at 1000 nM per kinase across 50 compound-kinase pairs predicted to be inactive by the two-stage pwkrr model. Red vertical dashed lines serve as references for the inactive threshold of 25% inhibition.
